# A gene expression panel for estimating age in males and females of the sleeping sickness vector *Glossina morsitans*

**DOI:** 10.1101/2021.05.12.443792

**Authors:** Eric R. Lucas, Alistair C. Darby, Stephen J. Torr, Martin J. Donnelly

## Abstract

Many vector-borne diseases are controlled by methods that kill the insect vectors responsible for disease transmission. Recording the age structure of vector populations provides information on mortality rates and vectorial capacity, and should form part of the detailed monitoring that occurs in the wake of control programmes, yet tools for obtaining estimates of individual age remain limited. We investigate the potential of using markers of gene expression to predict age in tsetse flies, which are the vectors of deadly and economically damaging African trypanosomiases. We use RNAseq to identify candidate expression markers, and test these markers using qPCR in laboratory-reared *Glossina morstians morsitans* of known age. Measuring the expression of six genes was sufficient to obtain a prediction of age with root mean squared error of less than 8 days, while just two genes were sufficient to classify flies into age categories of ≤ 15 and >15 days old. Further testing of these markers in field-caught samples and in other species will determine the accuracy of these markers in the field.

## 1 Introduction

Vector-borne diseases represent major threats to health and livelihood world-wide, being directly responsible for 680,000 deaths annually (Roth et al. 2018), as well as causing huge economic damage to livestock (Eisler et al. 2003, Shaw 2004). Control of the vectors that transmit these diseases is an integral tool for reducing disease burden (Wilson et al. 2020). The metric of success for these control programmes is a reduction in disease burden in the host population. However, when vector control is accompanied by other interventions such as screening and treating the host population for the disease, the contribution of vector control to the subsequent reduction of disease can be hard to determine (World Health Organization 2012). Conversely, while the impact on the vector population may not bear a simple relationship to disease burden, it is a direct outcome of vector control. Control efforts should thus be accompanied by detailed monitoring of the targeted vector populations, to estimate impact, to monitor population recovery and to understand the transmission dynamics of the disease. Mostly, monitoring currently relies on counting the number of vectors caught in sentinel traps, which can be greatly affected by trapping method, effort and efficacy, and may only partly reflect the ability of the vector population to transmit disease (Wilson et al. 2015).

One aspect of vector monitoring that has been particularly challenging is the quantification of the age-distribution (demographics) of natural populations (Caragata et al. 2011, Cook et al. 2006, Sikulu et al. 2010). Estimating vector age is important for two reasons. First, it can provide a measure of the effectiveness of vector control because increased adult mortality should lead to a younger population age structure. Importantly, this measure of control effectiveness is independent of catch size and trapping effort because only the distribution of age needs to be known. Second, in most cases, the probability that an individual vector is infectious for a given disease increases with age (Dye 1992, Woolhouse & Hargrove 1998). Before transmitting the disease, vectors first need to have taken an infected blood meal, and there is then typically a delay between acquisition of infection and onward transmission due to the need for the pathogen to replicate and/or mature. Age grading is therefore useful to determine the proportion of individuals old enough to transmit disease.

Tsetse flies (genus *Glossina*) are the vectors of Human African Trypanosomiasis (HAT, or sleeping sickness) and Animal African Trypanosomiasis (AAT, or nagana). HAT is, without treatment, a fatal disease endemic to sub-Saharan Africa (Franco et al. 2014), while AAT presents a major economic burden to rural communities by affecting livestock (Eisler et al. 2003). Being a disease primarily of animals and with reservoirs across multiple species, AAT cannot be controlled through treatment alone and is thus highly dependant on vector control (Holmes 2013). *G. morsitans morsitans* is a major vector of AAT in East and Southern Africa and can also transmit HAT (Dale et al. 1995). Catch rates of this species in the wake of vector control can be extremely low (Kgori et al. 2006, Vale et al. 1988, Van den Bossche 1997), making it particularly challenging to conduct ongoing monitoring of important populations. It is therefore all the more important to extract as much information as possible from the limited number of flies obtained.

As is the case for all insect vectors, a means to accurately determine the age of tsetse flies is a valuable but elusive goal, and current methods have many shortcomings. Laborious ovary dissections can be used to age females up to their fourth ovarian cycle (Hargrove 2012), but this technique requires specialist dissection skills and cannot be applied to males, despite males being at least as competent at transmission as females, and perhaps more so (Dale et al. 1995, Maudlin et al. 1990). Estimates of age based on wing damage (Hargrove 1990) or analysis of pteridines have also been used (Langley et al. 1988, Lehane & Hargrove 1988), but experience in practical applications has shown that measurements in the field vary enormously (for example in mosquitoes: (Lardeux et al. 2000, Penilla et al. 2002)) and cannot be used to reliably estimate age on an individual basis (Hargrove 2020).

Here we explore the value of using gene expression to estimate age in tsetse flies. This method has previously been tested in mosquitoes (Caragata et al. 2011, Cook et al. 2006), with encouraging results, but has yet to be applied in tsetse. We use laboratory-reared *G. morsitans* as a proof of concept, and show that measuring the expression of just six genes can estimate the age of both male and female tsetse flies with a root mean squared error of less than 8 days. We also trained models to classify tsetse into those younger or older than 15 days, since flies younger than 15 days are unlikely to harbour a mature trypanosome infection (Dale et al. 1995), and found that just two genes are sufficient for 95% accurate classification.

## 2 Methods

### 2.1 Sample collection and RNA extraction

*G. morsitans morsitans* individuals were collected from colonies maintained at the Liverpool School of Tropical Medicine. Colonies are kept in meshed boxes (cages) at 26°C ± 2 °C and 72 ± 4% humidity, with a 12hr light-dark photoperiod, and fed three times per week using defibrinated horse blood (TCS Biosciences Ltd., Buckingham, UK) provided through silicon-membrane feeders. Pupae are regularly collected and allowed to emerge to form new cages. Each fly cage contains flies which eclosed over a 2-3 day window, and thus the age of all flies in the cage are known to a precision of either 2 or 3 days. The ages reported here are the middle of the age range (eg: a fly aged 13-15 days or 13-16 days is reported as 14 days old). The age of the samples ranged from 2 to 62 days. While reproductive status of females was not measured precisely, we tried to include a range of physiological states (based on visual inspection of the size of the abdomen) within each age group, so that genes could be identified that are predictive of age in spite of variation caused by the ovarian cycle. Overall, 505 flies were collected (301 female and 204 male, Supplementary Data S1).

For sample collection, fly cages were briefly transferred to a cold room (4 °C) where flies to be collected were removed from the cage once quiescent and decapitated. Heads were placed into RNAlater and stored at −20 °C. In case repeated exposure to the cold room created alterations in gene expression, we minimised this exposure by never collecting flies from a given cage more than three times over the course of the experiment. No more than two flies were collected from a cage on a given day, for three reasons. Firstly, we wanted to make sure that flies were obtained from a range of different cages in order to avoid issues of results being confounded by cage of origin (such as an infection specific to one cage of flies). We therefore never obtained more than six flies from a single cage over the course of the experiment. Second, we wanted to minimise the time that samples spent at temperatures above −20 °C after death, limiting the number of samples that could be collected in a single sitting. Third, all flies were collected at the same approximate time of day (morning) to minimise gene expression variation due to circadian cycles (Rund et al. 2011), limiting the number of collections that could be performed on the same day.

RNA was extracted from individual fly heads. Single heads contain enough material for RNA sequencing and can easily be removed without the need for precise dissection, providing a quick and convenient tissue for sampling. We avoided the abdomen because of the important effect that sex and the ovarian cycle would have on gene expression in these tissues. RNA extractions were performed using PicoPure kits (Arcturus), increasing the volume of extraction buffer and alcohol to 120μl. cDNA libraries were prepared using SuperScript III Reverse Transcriptase (Invitrogen).

### 2.2 Sequencing

cDNA libraries from 22 male and 28 female individual flies ranging in age from 2 to 62 days post-eclosion (Fig. 1, Supplementary Data S1) were sent to the Liverpool Centre for Genomic Research (CGR) for 150bp paired-end sequencing on an Illumina HiSeq 4000 sequencer. Strand-specific library preparation was performed using NEBNext poly A selection and Ultra Directional RNA library preparation kits, producing an average of 23.8 million reads per sample. Reads were then trimmed as part of the CGR’s genomic pipeline using Cutadapt version 1.2.1 (Martin 2011) with option -O 3 to remove Illumina adapter sequences, and Sickle version 1.2 (https://github.com/najoshi/sickle/releases/tag/v1.2) with a minimum window quality score of 20. Reads shorter than 20 bp after trimming were removed and subsequently unpaired reads were excluded. Data were quality checked using FastQC (Andrews 2010) before analysis.

**Figure 1:**
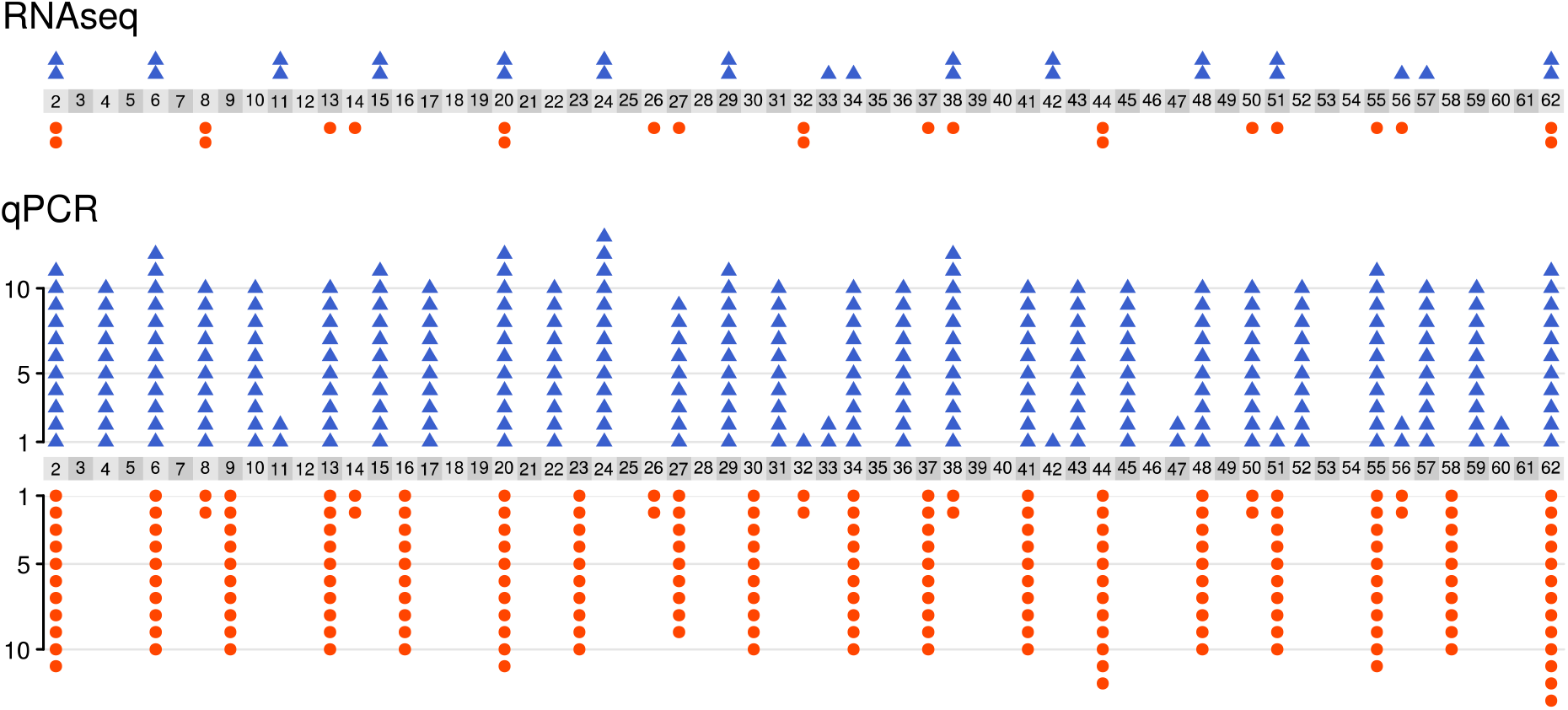
Number of samples used for RNAseq (top; total = 50) and qPCR (bottom; total = 498), split by age category (2 - 62 days old). Individual female and male flies shown as blue triangles and orange circles respectively.

### 2.3 RNAseq analysis

Trimmed reads were aligned to the GmorY1.9 genome using STAR aligner version 2.7.0 (Dobin et al. 2013) using the --quantMode GeneCounts option to obtain mapping counts for each gene.

Differential expression analysis was performed using the R package *EdgeR* (Robinson et al. 2010), with library size normalisation performed using Trimmed Mean of M-values (Robinson & Oshlack 2010) and dispersion calculated with trended and tag-wise estimates. Genes with fewer than 10 reads across all 50 samples were excluded from the analysis. All plotting figures show expression measured as reads per million reads (RPM) from normalised library sizes. Association of gene expression with age and sex was tested using generalised linear modelling (glm) implemented in *edgeR*, with age coded as a continuous variable and sex as a categorical variable. Preliminary analysis found little evidence of an important effect of the number of times a colony was exposed to the cold room on gene expression, but there was a significant effect of the number of days since flies had received a blood meal (Supplementary Data S2). We therefore controlled for days since receiving a blood meal by including it as a fixed continuous factor in the glm. False discovery rate control was set at 1% using the R package *fdrtool* (Klaus & Strimmer 2015).

Gene clustering analysis was performed with the *WGCNA* package in R (Langfelder & Horvath 2008), using the normalised read counts generated by e*dgeR* and keeping only the 5000 genes with the highest variance in expression. We used the hybrid module merging algorithm with a deep split value of 4, a minimum cluster size of 30 and a power parameter of 8, followed by module merging using the absolute value of the correlation coefficient between eigengenes as a distance matrix and a merging threshold of 0.2.

Prediction of age based on normalised read counts from the RNAseq data was performed using lasso regression implemented with the *glmnet* package in R (Friedman et al. 2010). As the aim was to find genes with consistently high predictive value for age, we explored a range of lasso parameters. This exploratory procedure is recorded in detail in the R script “02_lasso.r” provided on GitHub (https://github.com/EricRLucas/TsetseAgeMarkers).

### 2.4 Primer design and qPCR

Based on the results of the RNAseq analysis, 16 genes were short-listed to be tested as qPCR markers of age in *G. morsitans*, with two further genes being identified as suitable housekeeping genes for our purposes (i.e.: showed minimal variation in expression in the conditions included in our study and no evidence of association with age). Primers were designed for these genes based on the GmorY1.9 genome using NCBI Primer blast (Ye et al. 2012). Where possible, amplicons were designed to span exon junctions. Based on testing amplification efficiency using 1:3 serial dilutions, the 10 best primer pairs for age-predictive genes, and the two primer pairs for housekeeping genes, were kept for use in the study and applied to 499 samples (298 females and 201 males), including 44 of the samples used for RNAseq (the remaining 6 samples had too little cDNA left to be included in the qPCR study). One of the samples failed to produce a Ct value for several genes and was therefore excluded from subsequent analysis, leaving 498 samples (Fig. 1). All primers used in this study are listed in Supplementary Data S3.

qPCR was run on a AriaMX RealTime PCR instrument in a total volume of 20 μl, containing 10 μl of SYBR 2x MM, 1.2 μl of forward primer (5μM), 1.2 μl of reverse primer (5μM), 6.6 μl of nuclease-free water and 1 μl of genomic DNA. Reaction conditions: one cycle of 95°C (3 minutes), 40 cycles of 95°C (10 seconds) and 60°C (10 seconds), one cycle of 95°C (1 minute), 55°C (30 seconds) and 95°C (30 seconds, 5 seconds soak time).

Missing raw Ct values for age-predictive genes (where the signal never reached the threshold even after 40 cycles) were replaced with the maximum value of 40. ΔCt values were calculated using the mean Ct of the two housekeeping genes. Where Ct values were missing for either housekeeping gene, normalisation was impossible and the normalised aging gene value was recorded as missing (NA). All samples were run in two technical replicates and the final ΔCt was taken as the mean of the two replicates. Gene GMOY005321 consistently showed variable ΔCt values between technical replicates, possibly due to low expression of this gene, and these values were kept unchanged. For all other genes, any gene-sample combinations whose ΔCt differed by more than 1 between technical replicates were rerun for a third technical replicate, along with both housekeeping genes, providing a third ΔCt. In most cases, this third ΔCt was very close to one of the first two and very different from the other, indicating which of the first two technical replicates was wrong. The final ΔCt was thus taken as the mean of the third replicate and whichever of the first two replicates it was closest to.

### 2.5 Predicting tsetse age from qPCR data

Machine learning predictions of tsetse age from qPCR data were performed using the *caret* package in R (https://cran.r-project.org/package=caret). The ΔCt values for each of the 10 study genes were used as continuous predictor variables, and sex was included as a categorical predictor variable since some of the genes showed sex-dependent expression. Samples were randomly split into training set (75% of samples) and test set (25% of samples), stratified by sex and age to ensure equal representation of these two variables in the two sets. Due to rounding of sample numbers within each stratification layer, the final numbers in the train and test sets were 380 (76%) and 118 (24%) samples respectively. Model training was performed using three rounds of 10-fold cross-validation. For regression models, whose aim is to estimate age as a continuous variable, partial least squares regression (PLS), random forest and extreme gradient boosting (XGB) models were all trained on the data and their predictive accuracies compared. Categorical models were trained to categorise individuals into ≤ 15 and >15 days old. Simple decision tree, random forest and XGB models were compared for these categorical models.

The minimum number of expression markers (genes) required to obtain accurate predictions of age was determined by training the models with different numbers of loci. For each of the random forest and XGB models, the ten genes were ranked according to their variable importance in the full model training described above (sex was found to have a variable importance of 0 in both cases, and was therefore excluded from these models). The models were then trained with all ten genes, the top nine genes, the top eight genes, and so on. For each set of genes, 20 models were trained with a different random split of training and test sets, to account for stochastic variation in model accuracy.

All statistical analysis was conducted in R version 3.4.4 (R Core Team 2015). Analysis scripts, qPCR raw data and RNAseq read counts are available on GitHub (https://github.com/EricRLucas/TsetseAgeMarkers). Raw sequencing will be submitted to ENA shotgun sequencing archive upon final acceptance of the paper for publication.

## 3 Results

We collected 301 female and 204 male *G. morsitans* flies of known age from laboratory colonies, ranging in age from 2 to 62 days old. An initial RNAseq analysis of 28 female and 22 male samples showed that gene expression in these samples was primarily affected by age, rather than sex or days since last blood meal (Fig. 2, Supplementary Fig. S1), although this was primarily due to the strong changes in gene expression found during the first 15 days of life, with older individuals clustering primarily by sex (Supplementary Fig. S2).

**Figure 2:**
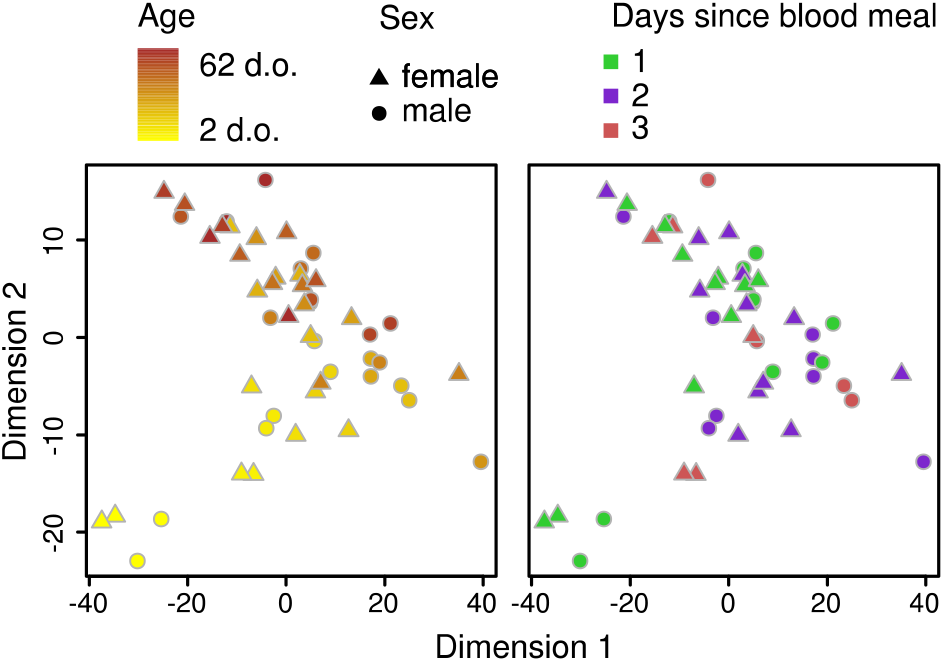
Gene expression clusters primarily by age. Principal component analysis of RNAseq data, coloured by age (left) or days since blood meal (right).

We identified a set of genes that was likely to provide strong age prediction by looking for genes that: 1. Were strongly correlated with age, or 2. consistently performed well in prediction of age using lasso regression and 3. where possible, belonged to different gene clusters as defined by weighted gene network clustering analysis. We particularly looked for genes showing strong expression changes in older individuals by identifying the genes most differentially expressed when considering only individuals older than 15 days, but even these showed relatively slight changes with age compared to some of the changes seen in the first 15 days of life (Fig. 3, Supplementary Fig. S3). Using our criteria, and after testing qPCR primer efficient, we manually picked 10 genes associated with age, and 2 genes with very little variation across samples to serve as housekeeping genes (Figs. 3 and 4).

**Figure 3:**
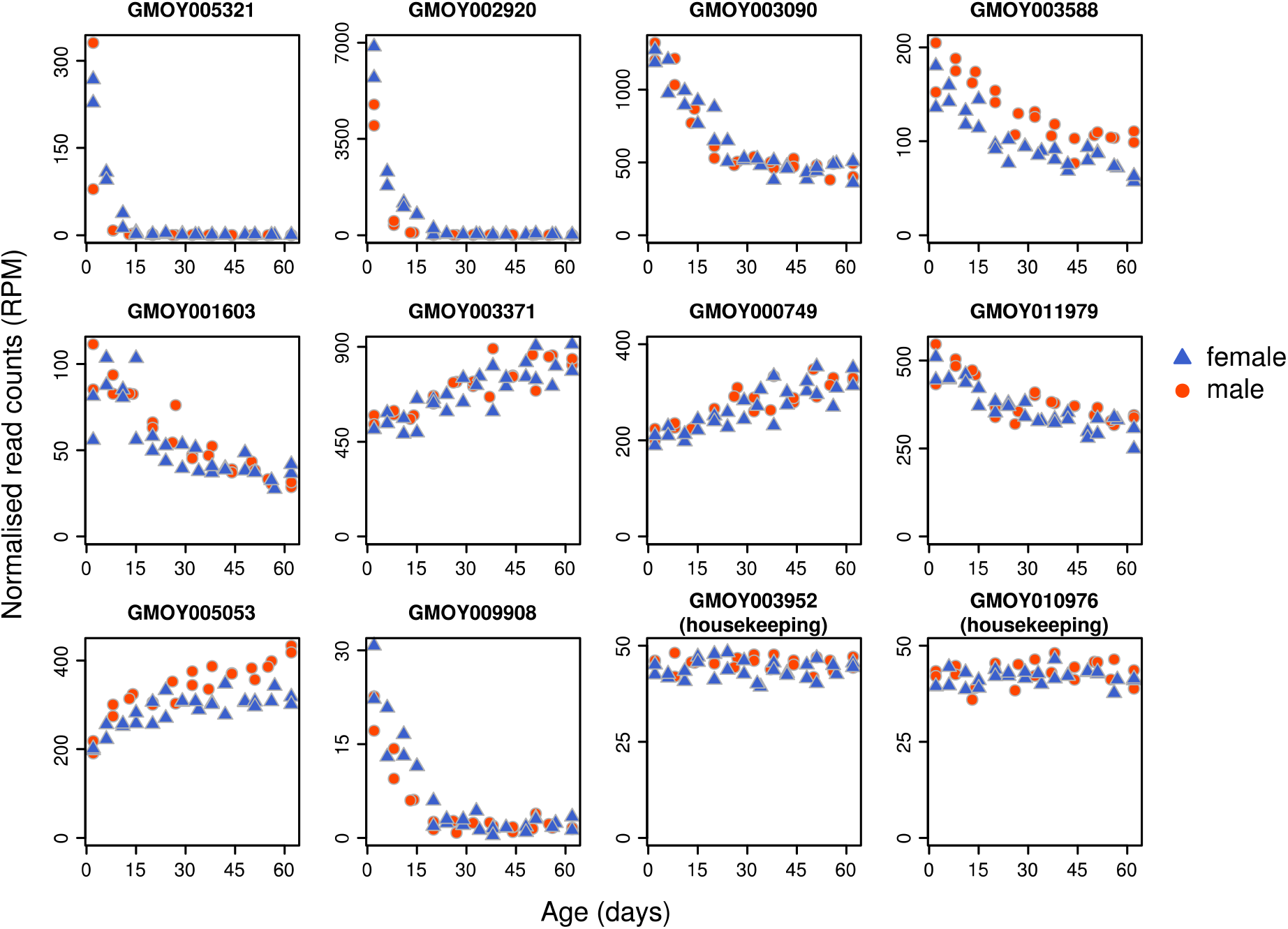
Expression of ten age-related genes and two housekeeping genes from RNAseq data, ordered according to the variable importance in the XGB model (Fig. 4). Very strong early-age expression changes in some genes (eg: GMOY005321, GMOY002920) allow good discrimination among young individuals, but show little change in later life. Genes with continuous changes (eg: GMOY003371, GMOY000749) are more gradual and offer more consistent, but less powerful, discrimination at all ages.

**Figure 4:**
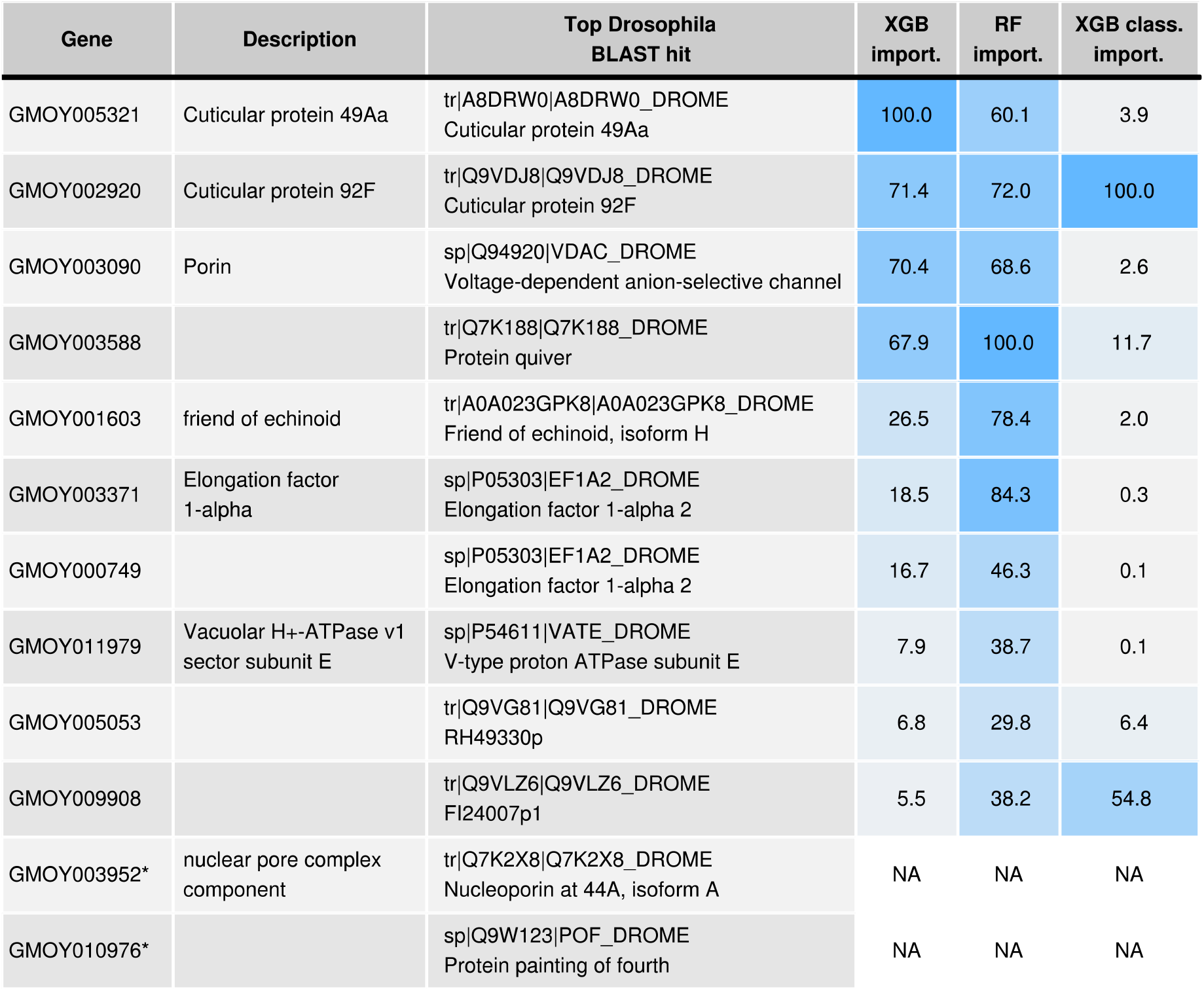
Ten age-related genes and two housekeeping genes (denoted with *) were used for qPCR analysis. Gene descriptions are taken from the Contig names in the GmorY1.9 proteome. Top Drosophila BLAST hits obtained by blasting the GmorY1.9 proteome against the D. melanogaster swissprot proteome. Variable importance of each gene shown for XGB, random forest (RF) and XGB classifier models trained with all predictor variables.

We obtained qPCR measurements of expression for these genes from 297 females and 201 males (Fig. 1). As expected, expression of all 10 age-related genes was strongly correlated with age (Supplementary Fig. S4) and with the RNAseq data (Supplementary Fig. S5). Principal component analysis of these age-related genes showed that age dominated the first principal component of the data. In particular, samples clustered strongly into those younger and older than 15 days (Supplementary Fig. S6)

The qPCR expression data produced strong overall predictions of age, with predictions being much more accurate in young flies (15 days or younger) compared to older flies. For regression models, PLS provided the poorest predictions of age, while random forest and XGB models performed equally well (Fig. 5, Supplementary Fig. S7). Taking the XGB model as an example, the overall root mean squared error (RMSE) for the final model was 6.74 days, but was 2.96 for individuals ≤ 15 days old. Variable importance for each gene in the random forest and XGB models are shown in Fig. 4. Training the model separately for males and females did not improve prediction accuracy (Supplementary Fig. S8).

**Figure 5:**
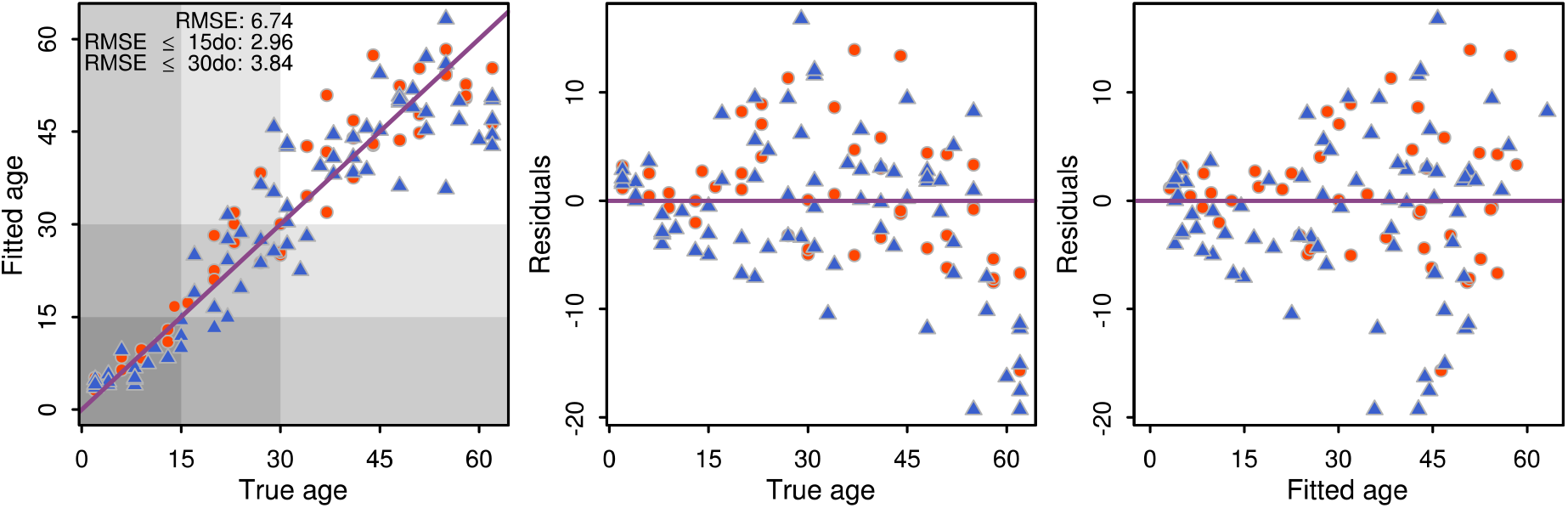
Prediction accuracy of the XGB model was highest (RMSE lowest) for individuals under 15 days old (2.59), and highest when all individuals were considered (6.81). Females are shown as blue triangles and males as orange circles. Purple line shows idealised perfect prediction.

Models also performed well at classifying samples into age categories of ≤ 15 and >15 days old (Supplementary Fig. S9). The XGB model performed best in this task, accurately classifying 117 out of 118 samples in the test set.

For both the random forest and XGB regression models, prediction accuracy showed little decrease when the variables of least importance were dropped from the models (Fig. 6). In both cases, accuracy remained comparable to that with all 10 genes when only 6 genes were included, with RMSE changing from 7.3 to 7.7 (random forest) or from 7.3 to 7.8 (XGB). In contrast, when moving to 5 genes instead of 6, RMSE changed from 7.7 to 8.4 (random forest) or from 7.8 to 9.3 (XGB). Interestingly, the same 6 genes proved to be sufficient for both model types (GMOY005321, GMOY002920, GMOY003090, GMOY003588, GMOY001603, GMOY003371). For the classification models, even fewer genes were needed (Fig. 6), with just two genes being sufficient for XGB classification accuracy consistently better than 95% (GMOY002920, GMOY009908).

**Figure 6:**
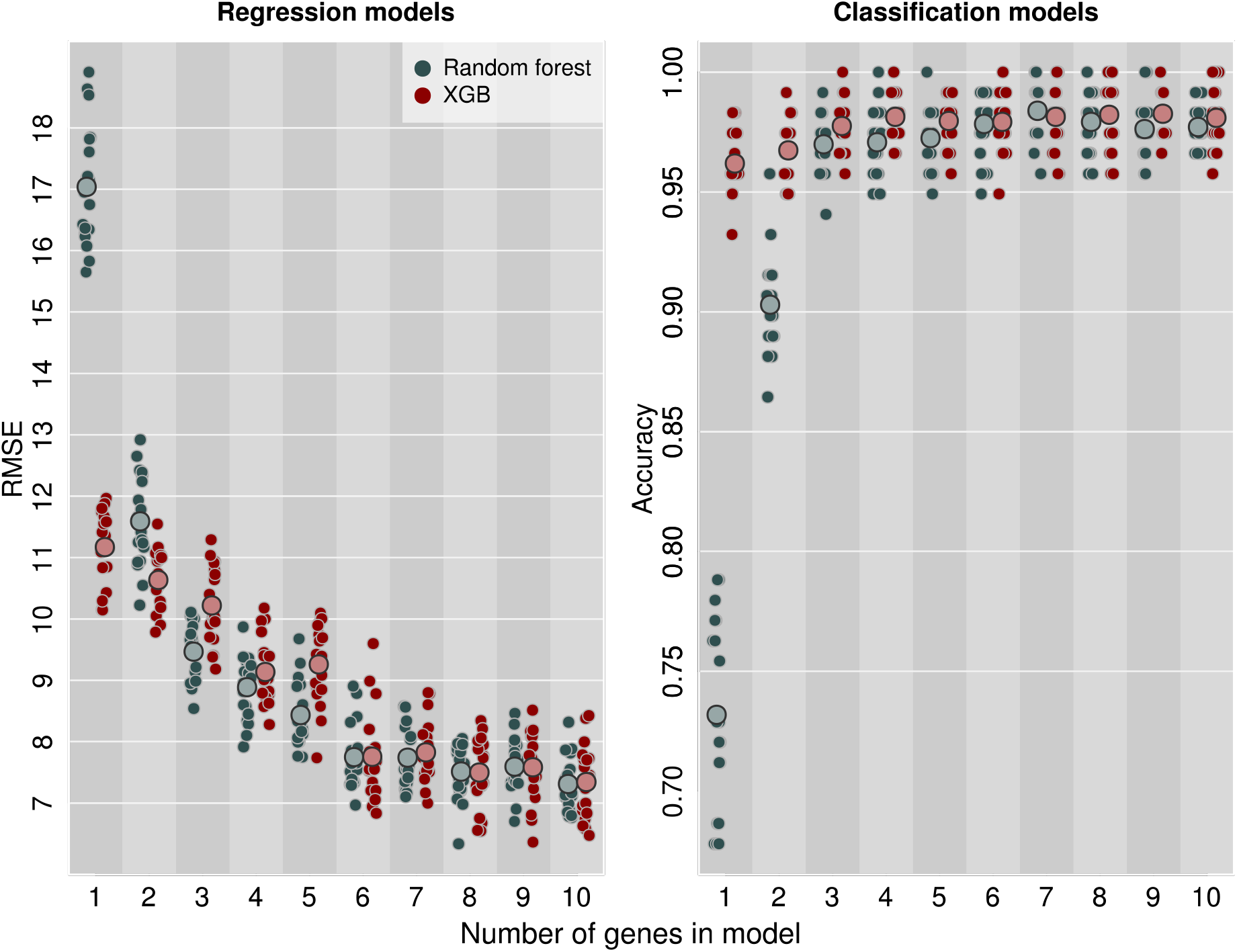
Predictive power of XGB and random forest models plateaus after the top 6 genes are included in the models (left). Accuracy of classification models plateaus after top 3 genes are included, with >95% accuracy achievable with only two genes (right). Small points show models run on independent test-train splits of the data (20 replicates per gene number); large points show the mean for each category. Points are jittered on the x axis to show overlapping data. bennett

## 4 Discussion

We have identified a set of gene expression markers that can be used to predict the age of *G. morsitans* tsetse flies in the laboratory. Importantly, this method can be applied to both males and females, providing accurate estimates of age in male tsetse. This is particularly important since not only do both male and female tsetse flies transmit trypanosomes, but males appear to be more likely to develop transmissible infections (Dale et al. 1995, Maudlin et al. 1990). Our genetic markers were also unaffected by time since an individual’s last blood meal, making them more robust for use on wild-caught individuals, where such factors cannot be controlled. Further work is nevertheless required to test the applicability of these markers in field conditions, since other environmental variables may still affect expression. For example, temperature and humidity were constant in our rearing conditions, and all samples were collected around the same time of day, leaving the possibility that these factors may yet influence the expression of our markers.

Like other methods for estimating the age of vectors, prediction accuracy decreases at older ages (Brei et al. 2004, Cook et al. 2006, Cook & Sinkins 2010, Gerade et al. 2004, Liebman et al. 2015, Penilla et al. 2002, Sikulu et al. 2010). In our data, this was because the change in expression with age was much greater in younger compared to older individuals, suggesting that the overall physiology of tsetse changes slowly after a certain life stage, and that there is thus little to detect that can be used for age grading. While we found genes that continued to change in older ages, the rate of change relative to the variance within age groups was not sufficient to achieve the same prediction accuracies as found in younger individuals. While it is likely that more accurate old-age predictions would be achievable using whole-transcriptome methods such as RNAseq, this is too costly to be applied at the scales required for training predictive models. In mosquitoes, spectroscopy-based methods used to estimate age initially suffered from a similar loss of precision at older ages (Liebman et al. 2015, Mayagaya et al. 2009, Sikulu et al. 2010, Sikulu-Lord et al. 2016), but recent studies using machine learning prediction methods have improved prediction accuracies (Lambert et al. 2018, Milali et al. 2019). Whether similar performance can be achieved with tsetse should be explored.

While we used ten genes in our study, we found that using only the six genes most predictive of age still provided high prediction accuracy, and only two genes were needed for classifying individuals into age groups of ≤ 15 and >15 days old. By removing four genes from the analysis, qPCR time and costs can be reduced by 1/3 (eight qPCR reactions per sample instead of twelve), while removing eight genes will reduce costs by 2/3. We thus suggest that further studies testing the applicability of these markers in the field restrict themselves to either six or two genes, depending on how precisely age needs to be estimated. Such studies are needed to determine the applicability of these markers in the field, but it would also be interesting to measure the expression of these genes in age-controlled samples of other species of tsetse to determine whether these markers have widespread applicability. Once the field applicability of these markers is confirmed, the technique can be rolled out in the context of monitoring of tsetse control campaigns by comparing the age distribution before and after interventions to confirm that a resulting shift in the population age distribution is observed. In particular, in the wake of a 100% effective campaign, no flies older than the start of the campaign should be found. The resulting data on age structure both before and after control campaigns can then also be used to inform epidemiological models of trypanosomiasis transmission.

In conclusion, our study provides a new method for estimating the age of tsetse flies which does not require specialist dissection skills and can be applied to males. The problem remains of finding methods for more accurately estimating age in older individuals. This may involve identifying senescent changes whose rate is steady and consistent enough to be generalisable to any individual in the population.

## Acknowledgements

We are grateful to Rob Leyland for assistance and tuition for the insectary work, and to Tom Churcher and Ben Lambert for valuable discussion on analytical approach. This work was supported by a Liverpool School of Tropical Medicine internal grant (Director’s Catalyst Fund) to ERL and a Medical Research Council, UK (MR/T001070/1) grant to MJD and ERL. SJT received support from the UK’s Biotechnology and Biological Sciences Research Council (grant numbers: BB/S01375X/1, BB/S00243X/1, BB/P005888/1) and the Bill and Melinda Gates Foundation (INV-001785, OPP1155293).

