## Supplementary Data S2 for "A gene expression panel for estimating age in males and females of the sleeping sickness vector *Glossina morsitans*"

### Electronic Supplementary Material

### Supplementary data S1

### Preliminary analysis of factors affecting gene expression

The effects of four factors on gene expression were studied, using the glmLRT function in *edgeR*. These factors were age, sex, number of previous exposures to the cold room ("box.uses" in the code below) and days since last blood feed ("days.since.feed"). The full script of this analysis is made available in the GitHub repository `https://github.com/EricRLucas/TsetseAgeMarkers` (file "analysis_day_and_cooling.r"), but in brief, the model and tests were coded as follows:

```
library(edgeR)
library(fdrtool)

design.matrix <- model.matrix(~age + sex + box.uses + days.since.feed)
read.counts.dgelist <- DGEList(counts = read.counts)
read.counts.dgelist <- calcNormFactors(read.counts.dgelist)
read.counts.dgelist <- estimateGLMTrendedDisp(read.counts.dgelist, design.matrix)
read.counts.dgelist <- estimateGLMTagwiseDisp(read.counts.dgelist, design.matrix)
read.counts.glm <- glmFit(read.counts.dgelist, design.matrix)

age.test <- glmLRT(read.counts.glm, coef = 'age')
age.ps <- age.test$table$PValue
age.fdr <- fdrtool(age.ps, statistic='pvalue')$qval

sex.test <- glmLRT(read.counts.glm, coef = 'sex')
sex.ps <- sex.test$table$PValue
sex.fdr <- fdrtool(sex.ps, statistic='pvalue')$qval

dsf.test <- glmLRT(read.counts.glm, coef = 'days.since.feed')
dsf.ps <- dsf.test$table$PValue
dsf.fdr <- fdrtool(dsf.ps, statistic='pvalue')$qval

uses.test <- glmLRT(read.counts.glm, coef = 'box.uses')
uses.ps <- uses.test$table$PValue
uses.fdr <- fdrtool(uses.ps, statistic='pvalue')$qval
```

Age and sex had the strongest effects on gene expression, followed by days since last blood meal (Fig. **SD1**). After false discovery rate control at 0.01, the number of genes significantly affect by age, sex and days since last blood meal were 2749, 2889 and 620 respectively (out of a total of 11,278 genes). No genes were significantly affected by number of previous exposures to the cold room. We therefore performed the main analysis with only age, sex and days since last blood meal in the model, that is:

```
design.matrix <- model.matrix(~age + sex + days.since.feed)
```

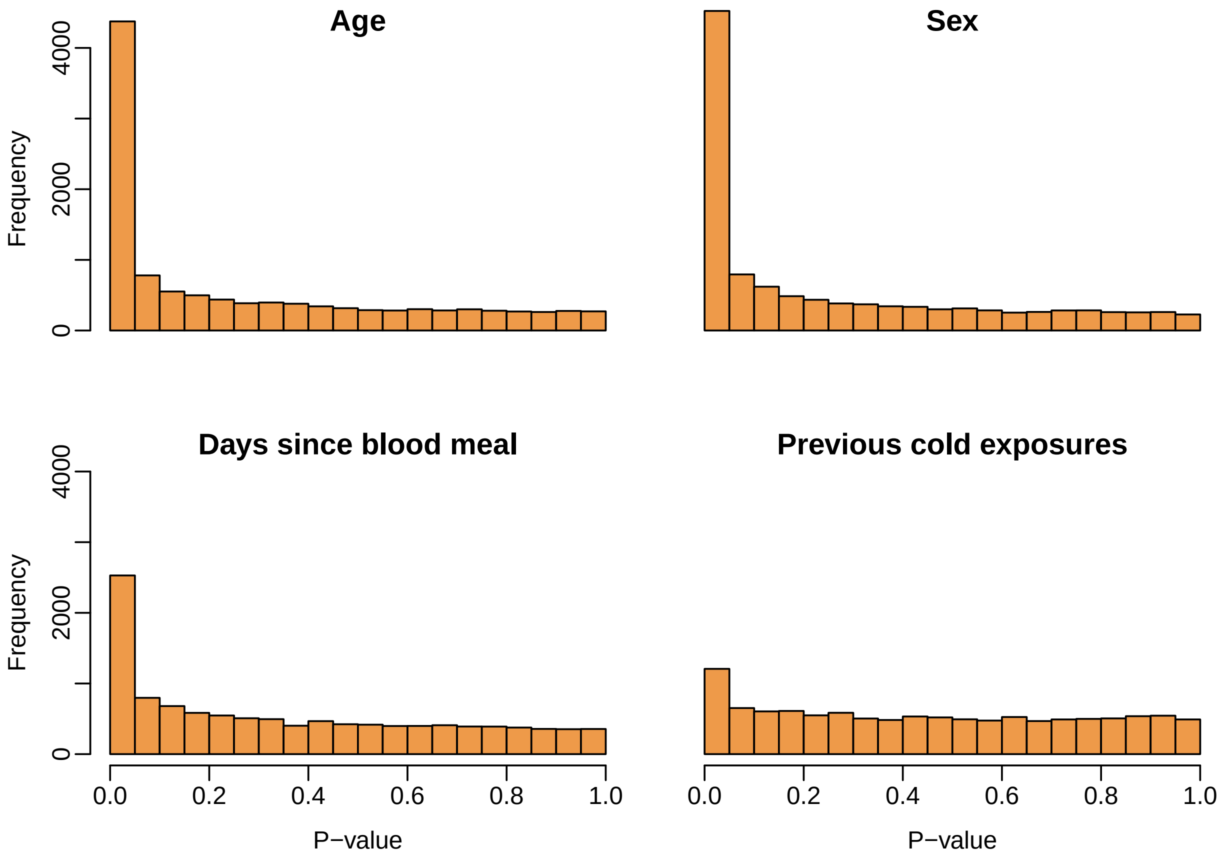


**Fig. SD1**: $P$-value histograms for the effects of age, sex, days since last blood meal and number of previous exposures to cold room. Factors with no effect on gene expression should approach a uniform $P$-value distribution. The number of truly differentially expressed genes in each bin can be roughly estimated by the height of the bin above the baseline defined by the height of the bins with $P$-values close to 1. Age and sex both had comparable effects on gene expression in terms of number of genes affected, followed by days since last blood meal. Number of previous exposures to the cold room had a negligible effect on expression.
