## Supplementary Figures for "A gene expression panel for estimating age in males and females of the sleeping sickness vector *Glossina morsitans*"

### Electronic Supplementary Material
### Supplementary figures and tables

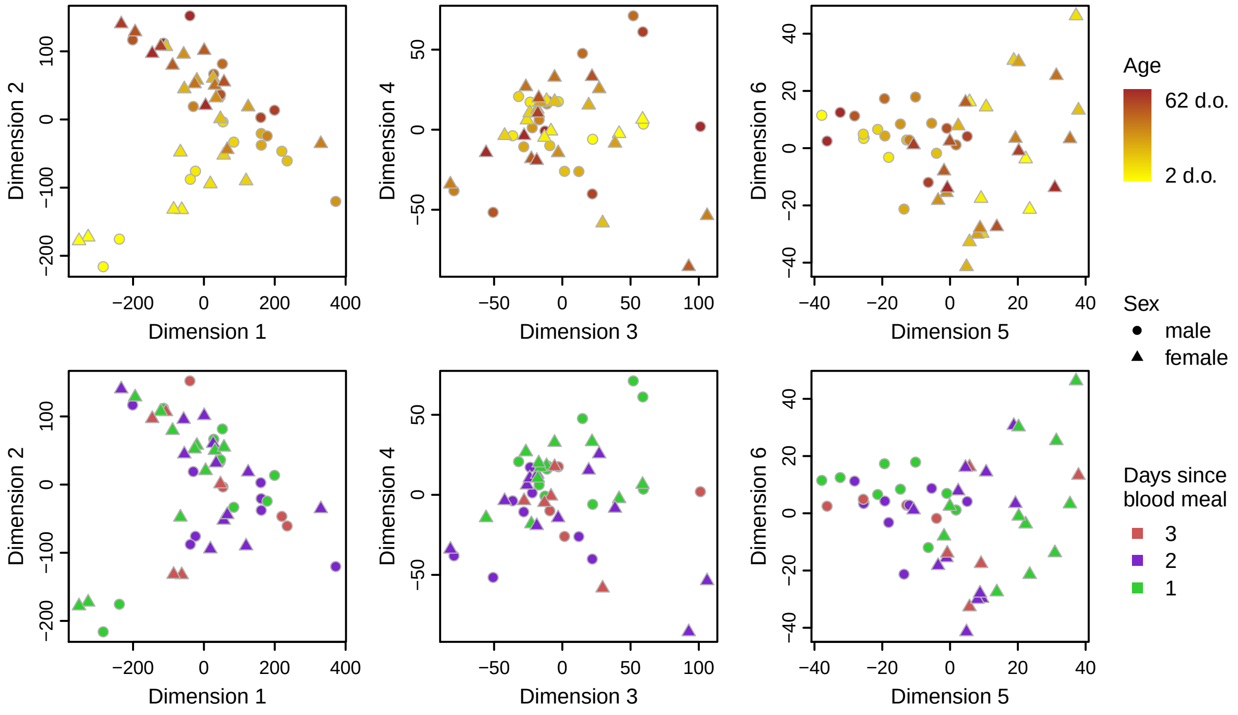


**Fig. S1**: Principle component analysis (PCA) of RNAseq data, coloured by age (top) or days since blood meal (bottom).

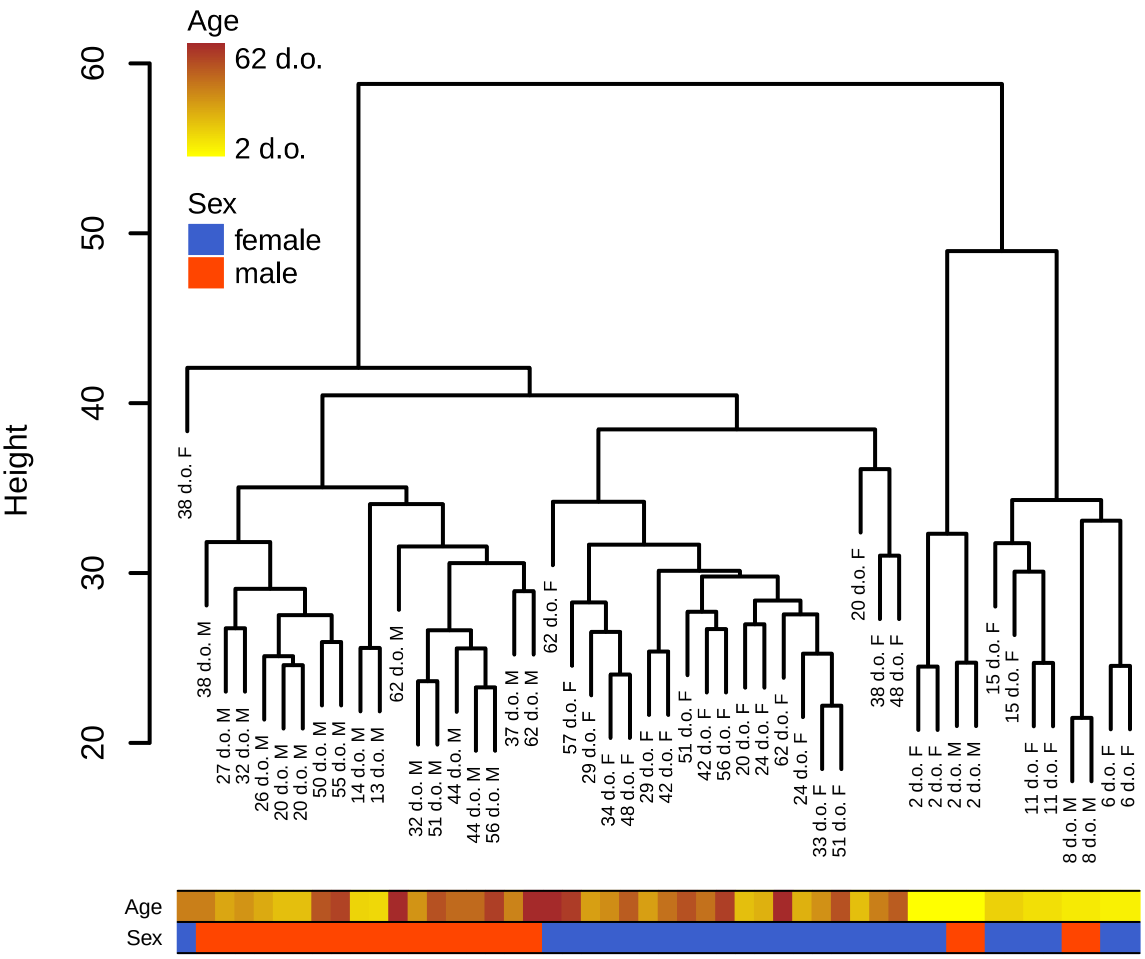


**Fig. S2**: Hierarchical clustering of samples from RNAseq data reveal that young individuals (<15 days old) cluster together, with older individuals clustering by sex.

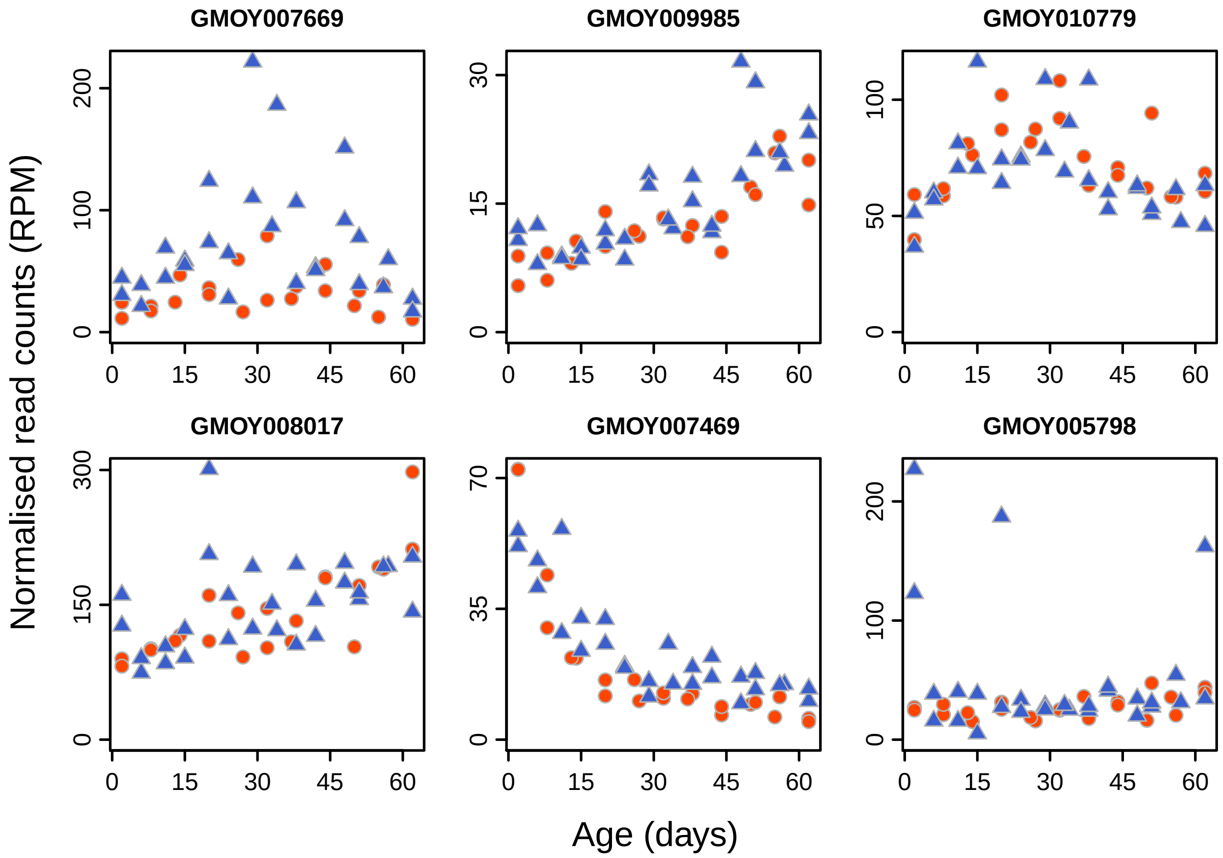


**Fig. S3**: Expression changes with age for the 6 genes most strongly differentially expressed by age when only individuals older than 15 days were included in the model. Females shown as blue triangles; males shown as orange circles.

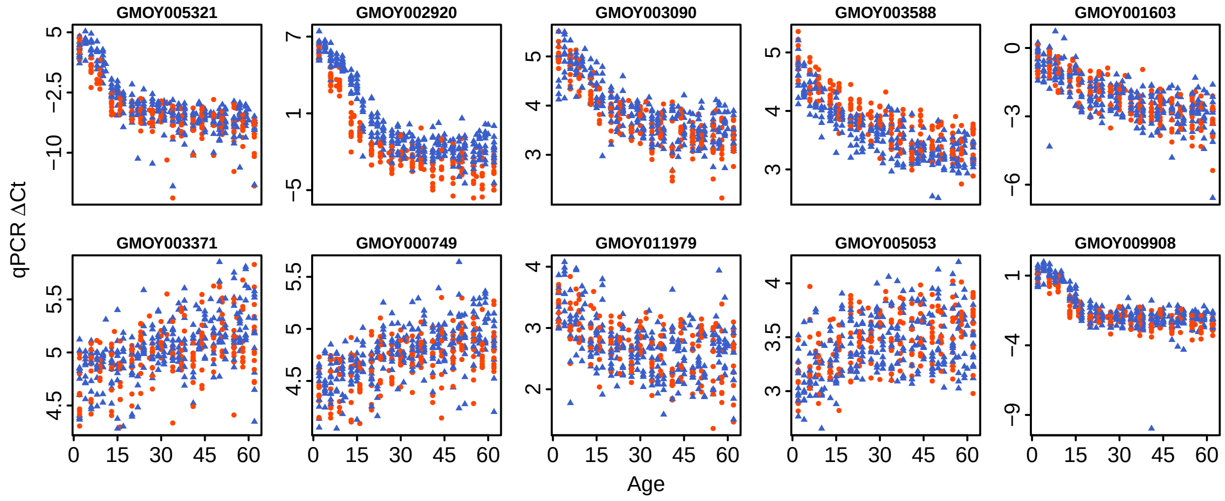


**Fig. S4**: Correlation of qPCR measures of gene expression against age for the ten genes chosen as age markers. Females shown as blue triangles; males shown as orange circles.

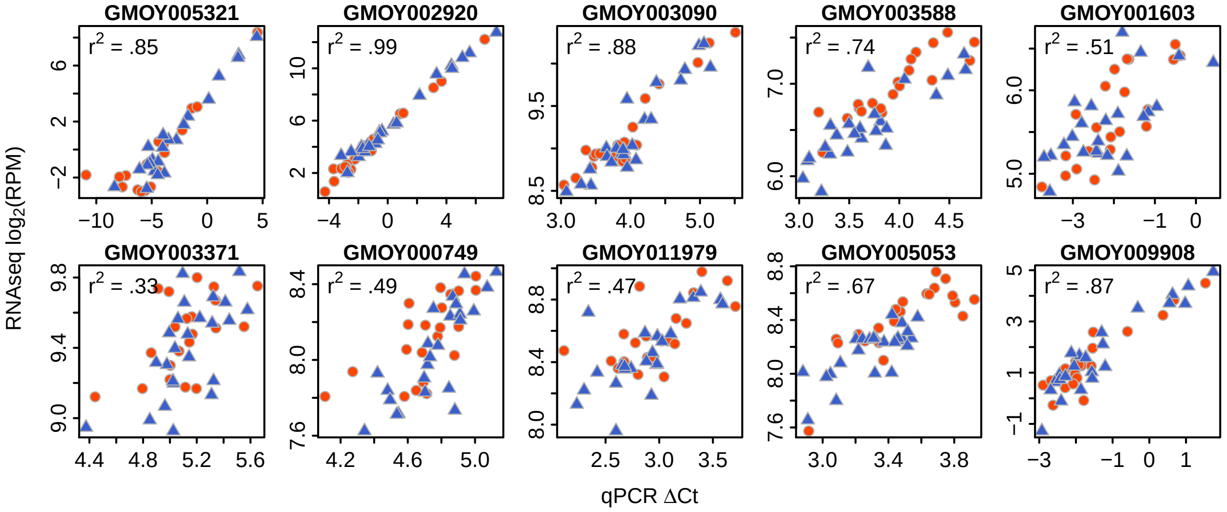


**Fig. S5**: Expression measured by qPCR and RNAseq were highly correlated in the samples in which both techniques were used. Expression by RNAseq is here measured as $\log_2(\text{RPM} + 0.1)$. RPM = reads per million. 0.1 added to avoid taking logs of 0. Females shown as blue triangles; males shown as orange circles.

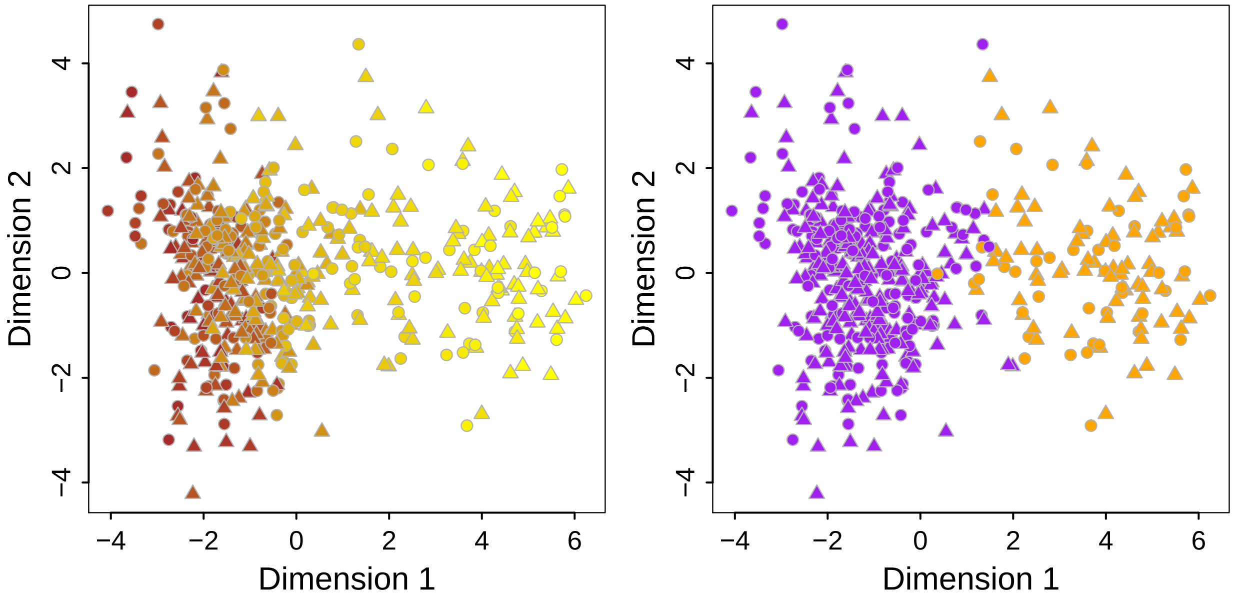


**Fig. S6**: PCA of samples based on qPCR measurements of expression of the 10 age-related genes. Left: colour-coded continuously from 2 days old(yellow) to 62 days old (brown). Right: colour-coded categorically into $\leq$ 15 days (orange) and $>$ 15 days (purple). Females shown as triangles; males shown as circles.

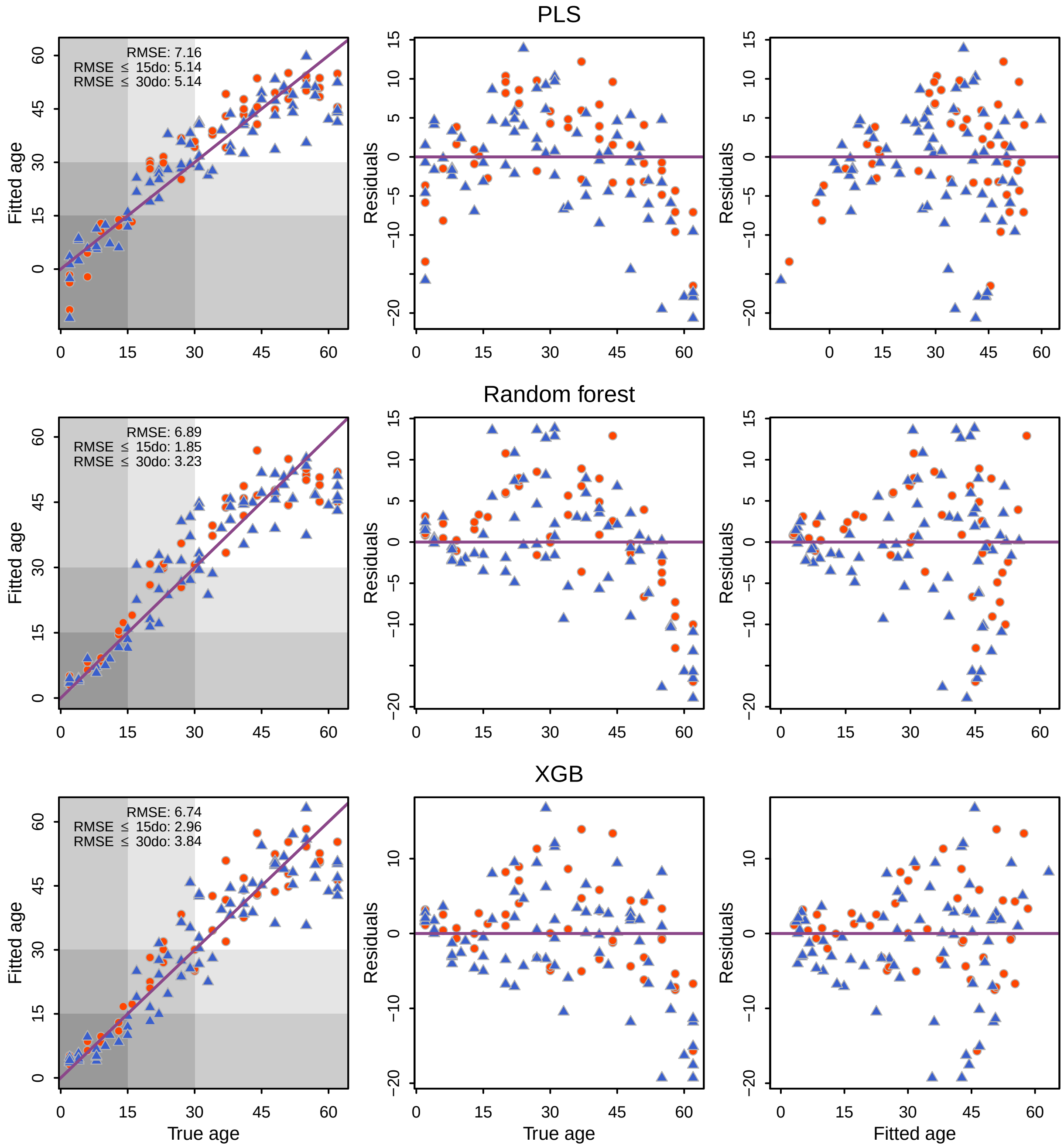


**Fig. S7**: Age prediction performance of PLS, random forest and XGB regression models. Females shown as blue triangles; males shown as orange circles.

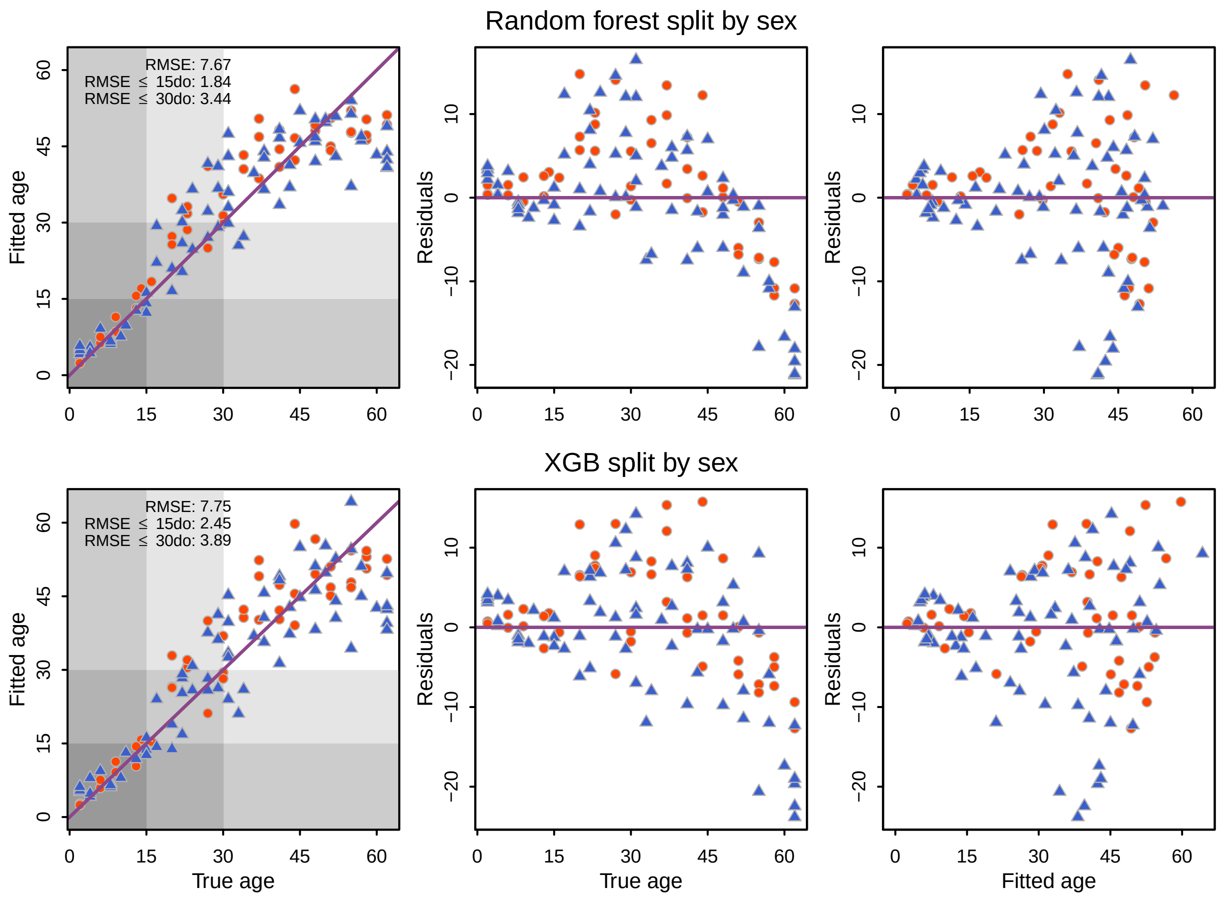


**Fig. S8**: Age prediction performance of random forest and XGB regression models trained separately on females and males. Females shown as blue triangles; males shown as orange circles. Purple line shows idealised perfect precdiction.

| | | *Decision tree prediction* | | *Random forest prediction* | | *XGB prediction* | |
|---|---|---|---|---|---|---|---|
| | | **Young** | **Old** | **Young** | **Old** | **Young** | **Old** |
| *True age* | **Young** | 25 | 3 | 27 | 1 | 28 | 0 |
| | **Old** | 0 | 90 | 1 | 89 | 1 | 89 |

**Fig. S9**: Accuracy at classifying samples into age groups of $\leq 15$ and $> 15$ days old was 99% for the XGB classification model, 98% for the random forest and 97% for the decision tree.
